# Formation of novel PRDM9 allele by indel events as possible trigger for tarsier-anthropoid split

**DOI:** 10.1101/047803

**Authors:** Sacha Heerschop, Hans Zischler, Stefan Merker, Dyah Perwitasari-Farajallah, Christine Driller

## Abstract

*PRDM9* is currently the sole speciation gene found in vertebrates causing hybrid sterility probably due to incompatible alleles. Its role in defining the double strand break loci during the meiotic prophase I is crucial for proper chromosome segregation. Therefore, the rapid turnover of the loci determining zinc finger array seems to be causative for incompatibilities. We here investigated the zinc finger domain-containing exon of *PRDM9* in 23 tarsiers. Tarsiers, the most basal extant haplorhine primates, exhibit two frameshifting indels at the 5’-end of the array. The first mutation event interrupts the reading frame and function while the second compensates both. The fixation of this peculiar allele variant in tarsiers led to hypothesize that de‐ and reactivation of the zinc finger domain drove the speciation in early haplorhine primates. Moreover, the high allelic diversity within *Tarsius* point to multiple effects of genetic drift reflecting their phylogeographic history since the Miocene.

## Introduction

Meiosis is a fundamental process to generate hereditary variability, thus being a driving force in evolutionary biology. Meiotic abruption is a cause for hybrid sterility which is a postzygotic barrier leading to speciation^1,2^. The question now increasingly being asked is therefore what type of genetic modification underlies species formation. In vertebrates *PRDM9* is, at present, the top candidate gene associated with hybrid sterility^1,3^. The gene is exclusively expressed by germ cells in the meiotic prophase I and encodes a zinc finger protein that specifies hotspots of recombination^4‒6^. The zinc finger domain evolves rapidly, both as to the number and sequence of repetitive motifs resulting in altered DNA-binding specificity, which in turn may lead to genetic incompatibilities promoting species divergence^3^. A recently published study on the evolutionary dynamics of the primate *PRDM9* zinc finger array revealed taxon-specific alleles across 18 haplorhine species, further strengthening its role in speciation processes^7^. However, tarsiers, the sole extant representatives of non-anthropoid haplorhine primates^8^, were not included. As the deepest offshoot within haplorhini they have a long independent history, possibly covering about 80 million years of primate evolution^9^. Fossils of *Tarsius* are scarcely reported and restricted to the middle Eocene and middle Miocene in Asia^10,11^. Assuming the fossil families of Omomyidae and Archicebidae as primitive tarsiiforms, extinct ancestors of tarsiers were widespread holarctic Eocene species^12^ and occurred as far back as 55 Mya^13^. Extant tarsiers, however, are endemic to insular Southeast Asia, where they fall into the three geographically and evolutionary distinct lineages of Western, Philippine, and Sulawesi tarsiers, with the latter constituting the most species-rich clade^14,15^. Even considering only the radiation of this ancient primate lineage in the Malay-Archipelago since the Miocene, allopatric speciation both between and within extant tarsier clades has left traceable molecular signatures of biogeographic events^16‒19^.

Due to their independent evolution within haplorhine primates and their marked diversification, especially evident over the last 2.5 million years on the Indonesian island of Sulawesi^19^, tarsiers represent the unique opportunity for exploring the evolutionary dynamics of the *PRDM9* gene and its role in speciation processes within primates and tarsiers in particular. We therefore sequenced the *PRDM9* exon encoding the zinc finger array of 23 tarsier individuals, including 21 specimen from Sulawesi and two non-Sulawesi specimen, one each representing the Western and the Philippine tarsier clade. Based on these data we investigated the functionality and allelic diversity of the zinc finger domain in tarsiers. We further tested the hypothesis of potentially postzygotic reproductive barriers among allopatric and parapatric species. In addition we inferred evolutionary relationships between zinc fingers of anthropoid primates^7^ and tarsiers, and discussed the possible function of *PRDM9* as driving force behind the anthropoid-tarsiiform split and the divergence of the tarsiiform lineage.

## Results

### Tarsier PRDM9 sequence characterization

The examined exon of Western, Philippine and Sulawesi tarsiers was homologous to exon 11 of the human PRDM9 gene and was about 1037-1793 bp long depending on the number of zinc finger repeats. It could be divided into two parts, the 5’ sequence and the 3’ C2H2 zinc finger array. Structural and functional properties of the zinc finger protein encoding exon are shown in figure 1.

**Figure 1:**
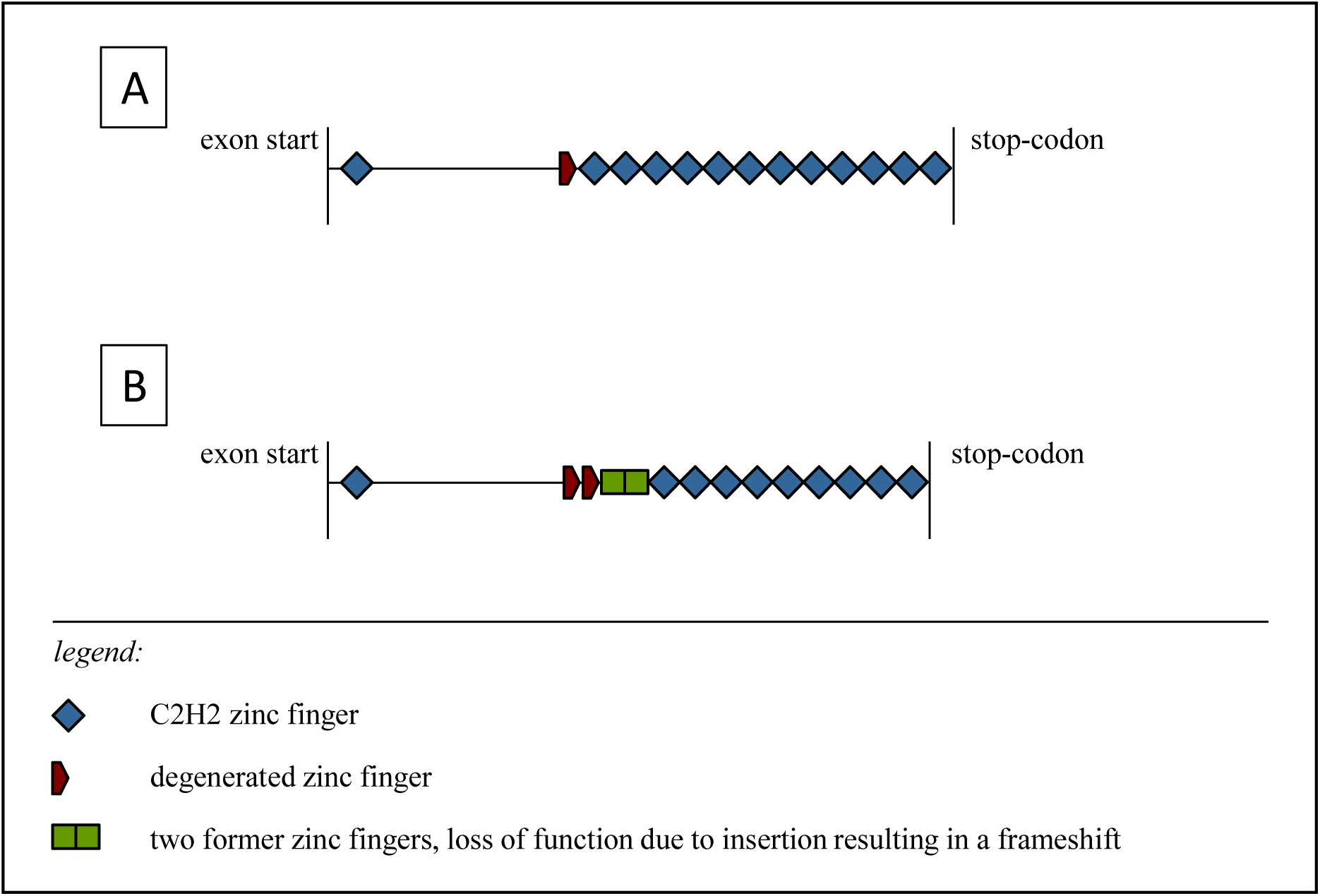
Schematic example of the *PRDM9* exon with the C-terminal zinc finger array of [A] primates and rodents and of [B] tarsiers. The comparison between the two exons shows the impact of the insertion and deletion on the zinc finger array, i.e. the loss of three functional C2H2 zinc fingers.

In contrast to the variable zinc finger domain the 5’ sequence of the exon was more conserved. This part of the sequence contained a C2H2 zinc finger at the beginning of the exon and four presumably former functional zinc fingers. The second and third zinc fingers were both degenerated by the loss of a zinc ion binding cysteine ligand, see figure 2, which is in principal not uncommon in zinc finger arrays^20^. The sequence of the third zinc finger was also interrupted by a 2 bp-insertion that caused a framing error affecting the next two zinc fingers. A deletion in the second altered zinc finger, which would be the fifth overall, restored the reading frame. The array of classical 84 bp C2H2 zinc fingers started with the end of this fifth zinc finger (Fig. 1, Fig. 2).

**Figure 2:**
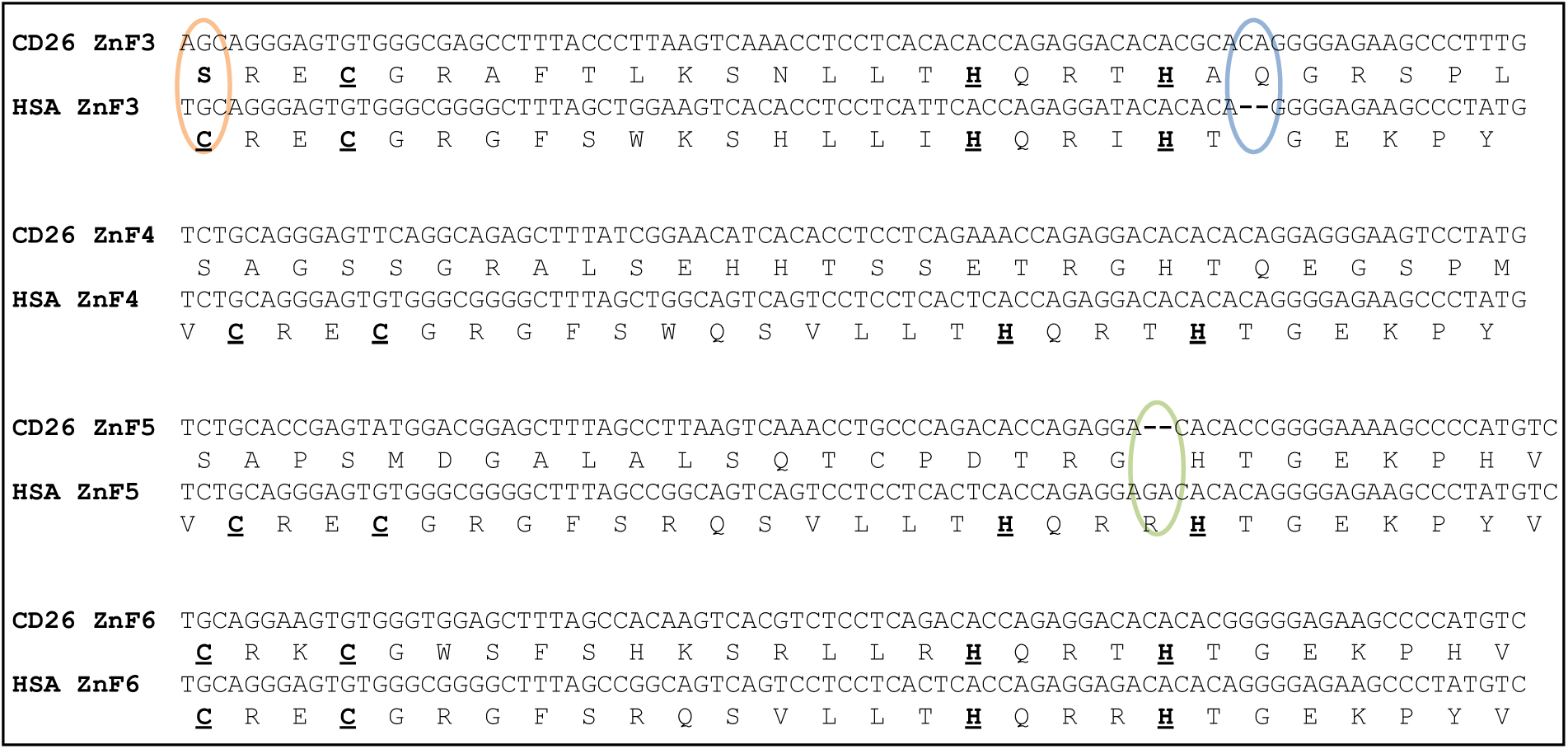
Frameshift mutations in tarsier *PRDM9*. Shown are aligned nucleic acid and peptide sequences comprising zinc fingers 3-6 (ZnF3-ZnF6) of *Homo sapiens* (HSA) and *Tarsius* (represented by a Sulawesian individual, CD26) and illustrating the frameshifting nature of insertion (blue) and deletion (green) mutations. Note the nonsynonymous substitution in the very first triplet which turns cysteine to serine (orange) and degrades ZnF3 of the tarsier. Amino acid sequences of human ZnF3-ZnF6 and ZnF6 of the tarsier show the classical C2H2 zinc finger motif (indicated by bold and underlined amino acids).

Indel events are quite rare in coding regions due to purifying selection^21,22^, especially if their length is not a multiple of three^23^. While the insertion results in a different amino acid sequence and may alter protein function, the deletion alone generates premature stop codons which lead to a faster pseudogenization respectively gene loss^24^. We therefore consider it likely that the insertion predates the deletion mutation which then in turn acted as a compensatory mutation. Compensatory mutations are twice as common as reversions to the original state and are often found in close proximity of the initial mutation^25,26^ being consistent with our findings. As only both indels jointly restore protein integrity we further assume that the temporal offset between indel mutations resulted in a temporary loss or limited protein function.

### Allelic variation of the zinc finger array in tarsiers

The alleles are defined on amino acid level, i.e. synonymous nucleotide changes are excluded, and are restricted to the C2H2 array. We found 28 alleles in 23 individuals with 15 individuals being heterozygous, (Table 1, Fig. 3).

**Table 1:** Allelic diversity of the zinc finger array in tarsiers.

| Individuals |  | Origin |  | Allele |  | Number of C2H2-ZnFs |  |
| --- | --- | --- | --- | --- | --- | --- | --- |
| Species | Sample ID | Biogeographic Region | Sample site | 1 | 2 | Allele 1 | Allele 2 |
| *Tarsius syrichta* | TSY | Philippines | N/A | 1 |  | 3 |  |
| *Tarsius bancanus* | TBA | Sundaland | N/A | 2 | 3 | 8 | 7 |
| *Tarsius dentatus* | T111 | Wallacea | Laone<br>Central Sulawesi | 14 | 15 | 9 | 8 |
| *Tarsius lariang* | T47 | Wallacea | Koja<br>Central Sulawesi | 16 | 17 | 10 | 7 |
| *Tarsius sp.* | CD02 | Wallacea | Ogatemuku<br>North-Sulawesi | 10 | 12 | 9 | 10 |
| *Tarsius sp.* | CD05 |  |  | 11 | 13 | 9 | 9 |
| *Tarsius sp.* | CD10 |  |  | 10 | 13 | 9 | 9 |
| *Tarsius dentatus* | CD14 | Wallacea | Korosule<br>Central Sulawesi | 18 | 21 | 6 | 7 |
| *Tarsius dentatus* | CD16 |  |  | 19 | 20 | 8 | 9 |
| *Tarsius dentatus* | CD17 |  |  | 18 | 22 | 6 | 6 |
| *Tarsius dentatus* | CD19 | Wallacea | Luwuk<br>Central Sulawesi | 14 |  | 9 |  |
| *Tarsius dentatus* | CD24 |  |  | 14 |  | 9 |  |
| *Tarsius dentatus* | CD26 |  |  | 14 |  | 9 |  |
| *Tarsius dentatus* | CD27 |  |  | 14 |  | 9 |  |
| *Tarsius sp.* | CD34 | Wallacea | Labanu<br>North-Sulawesi | 9 |  | 6 |  |
| *Tarsius sp.* | CD36 |  |  | 7 | 8 | 10 | 10 |
| *Tarsius sp.* | CD41 | Wallacea | Kendari<br>South-Sulawesi | 26 | 28 | 7 | 12 |
| *Tarsius sp.* | CD43 |  |  | 25 | 27 | 10 | 7 |
| *Tarsius sp.* | CD46 | Wallacea | Duasaudara<br>North-Sulawesi | 5 |  | 10 |  |
| *Tarsius sp.* | CD48 |  |  | 6 |  | 10 |  |
| *Tarsius sp.* | CD51 |  |  | 4 | 5 | 10 | 10 |
| *Tarsius fuscus* | CD64 | Wallacea | Bantimurung<br>South-Sulawesi | 23 | 24 | 6 | 11 |
| *Tarsius fuscus* | CD65 |  |  | 23 | 24 | 6 | 11 |

Each individual is listed with its species, sample ID, origin, allele identification and the number of C2H2 zinc fingers per allele. TSY: *Tarsius syrichta*; TBA.: *Tarsius bancanus*; *Tarsius sp.*: Taxon yet unclassified and currently affiliated with the *Tarsius tarsier*-population<sup>15</sup>.

The 25 alleles exclusively found in Sulawesi tarsiers show similarities, most often within populations (see figure 7 for sample sites). They mainly differ in the number of zinc fingers but hardly in their sequence like the alleles 9 and 10. Alleles 4, 5 and 6 have each ten zinc fingers with minor changes in the amino acid sequence (Fig. 3).

**Figure 3:**
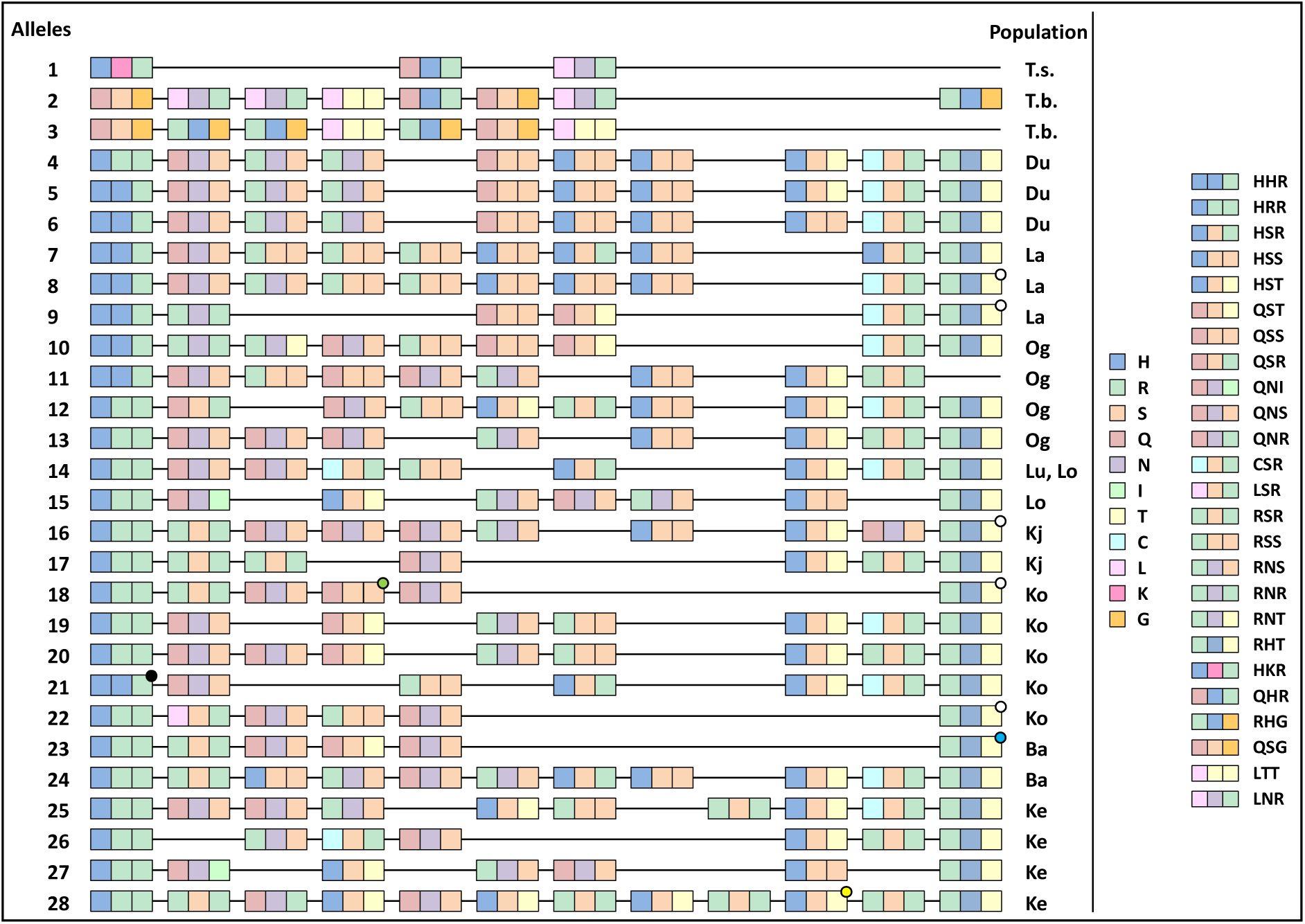
Alignment of C2H2 zinc finger alleles. The numbers identifying the alleles are identical to those in table 1. Each zinc finger is depicted with a triplet composed of its binding amino acids (-1, 3, 6). Each amino acid is marked with a distinct colour. Dots hint towards one or more nonsynonymous substitutions apart from the shown amino acids. Abbreviations: *T.s., Tarsius syrichta; T.b., Tarsius bancanus*; Du, Duasaudara; La, Labanu; Og, Ogatemuku; Lu, Luwuk; Lo, Laone; Kj, Koja; Ko, Korosule; Ba, Bantimurung; Ke, Kendari.

Only allele 14 is shared between two Sulawesian populations (Luwuk and Laone), both belonging to *Tarsius dentatus*.

There are two types of 5’-most zinc fingers differing in one binding amino acid (HHR, HRR). One variant can be found on the northern peninsula (Duasaudara, Labanu, Ogatemuku) and in Korosule. The other is specific to populations inhabiting the central, eastern and southern parts of Sulawesi, with the exception of one copy also occurring in north-eastern Sulawesi. The 3’-most zinc fingers are, with one exception, identical regarding the three binding amino acids RHT (Fig 3). Five of them show a nonsynonymous substitution which, however, does not affect the DNA-binding sites. They were observed in the populations of Labanu, Korosule and Koja (*Tarsius lariang*).

Three out of 19 zinc fingers with key codon positions QSR, LSR, and RNT are unique. Some are only found in few populations like the motifs specifying the binding triplet QNI or RNR and being restricted to Laone (*T. dentatus*) and Kendari, or Labanu and Ogatemuku, respectively.

Alleles of *T. bancanus* and *T. syrichta* are species-specific encoding zinc finger motifs not present in Sulawesi tarsiers. Despite their long geographic isolation they still share two zinc fingers (Fig. 3). Both alleles of the Western tarsier end with a zinc finger degenerated by the exchange of the second zinc binding histidine ligand with asparagine. With only three repeats the zinc finger domain of the homozygous Philippine specimen is unusually short, but according to Stubbs *et al*^21^ capable to bind DNA.

### Phylogenic analysis

To estimate the phylogenetic affiliations of the sequenced tarsier zinc fingers within the haplorhine divergence we reconstructed a phylogeny with *Microcebus murinus* as strepsirhine outgroup. Including all first degenerated and C2H2 zinc fingers (Fig. 4A) it completes the results of Schwartz *et al*^7^. from a haplorhine perspective. The basal trifurcation separates degenerated zinc fingers of tarsiers, degenerated zinc fingers of anthropoid primates and all C2H2 zinc fingers with node support values above 0.9 (Fig. 4A). Within the C2H2 cluster tarsier zinc fingers form a well-supported monophyletic group. Within this group 5’ most and 3’ most C2H2 zinc fingers are clustered reflecting the biogeography of the individuals tested. Apart from these findings our tree topologies (Fig. 4) are congruent with those obtained for anthropoids as presented by Schwartz *et al*.^7^.

**Figure 4:**
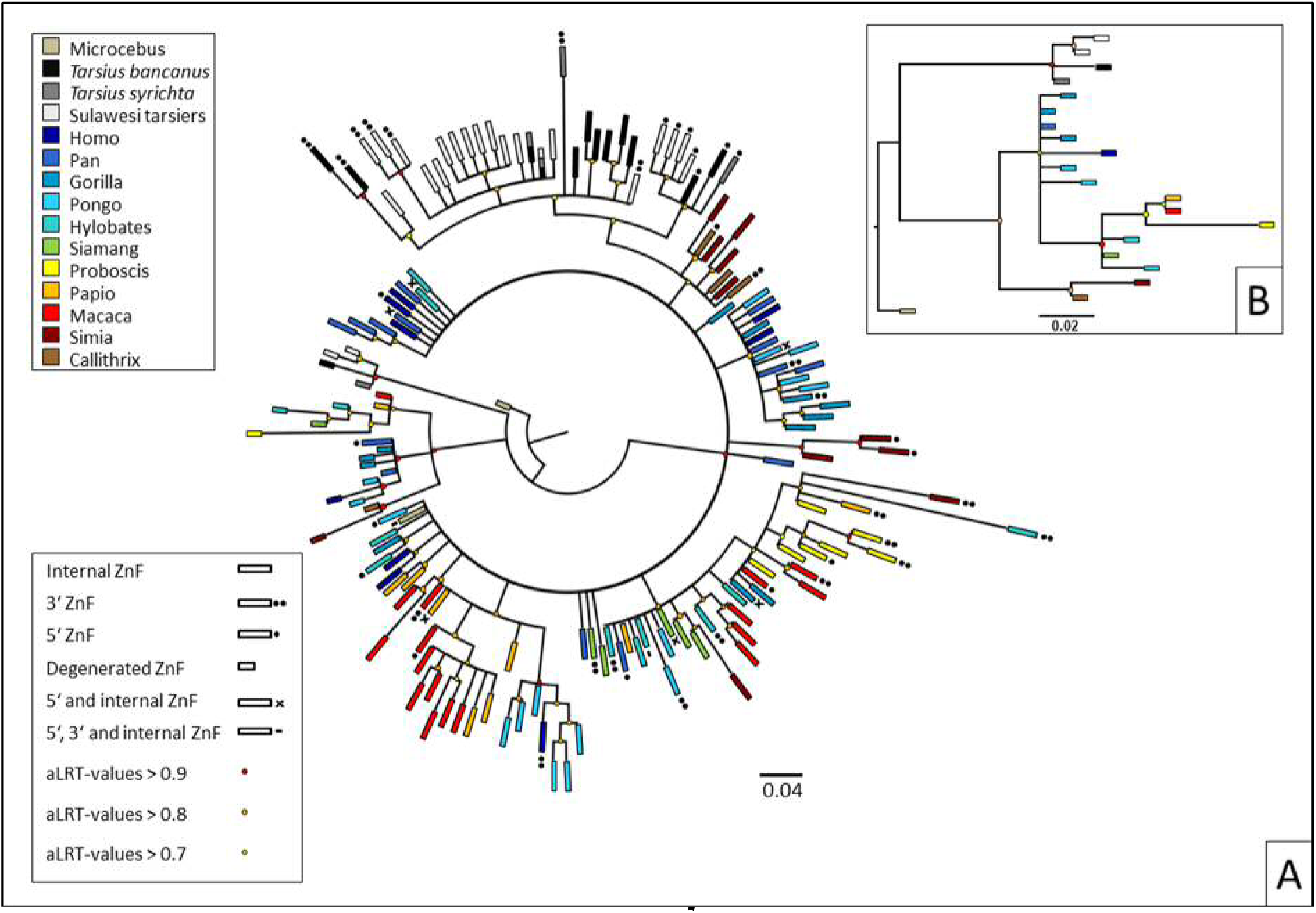
Phylogeny reconstructions based on Schwartz *et al*^7^, including tarsier zinc finger. [A] shows a tree topology including the first degenerated zinc finger and all C2H2 zinc fingers of the array. [B] depicts only the first degenerated zinc finger. The genera can be identified by colour, different kinds of or particular zinc fingers can be distinguished by their form, see key. Branches with aLRT-values above 0.9, 0.8 and 0.7 are indicated with a red, orange and yellow dot, respectively. The three main branches towards tarsier degenerated zinc fingers, anthropoid degenerated zinc fingers and towards C2H2 zinc finger show the highest aLRT-values of 0.992, 0.966 and 0.98, respectively.

The second phylogeny (Fig. 4B) is based exclusively on first degenerated zinc fingers. Here again tarsiers constitute a monophyletic group basal to the anthropoid clade. Within *Tarsius* Sulawesi tarsiers are monophyletic with respect to Western and Philippine tarsiers.

### Positive and negative selected sites

Zinc finger genes are known to alter quickly in their zinc finger array and in particular at the DNA-binding codons (positions −1, 3 and 6 relative to the a-helix) which show signatures of positive selection^28‒30^. The *PRDM9* zinc finger domain is no exception as shown by positively selected DNA-binding amino acids in rodents and primates^3,31,32^. The high mutation rates at codon positions directing the interaction of PRDM9 with DNA is deemed responsible for the formation of new hotspots and thus antagonizing hotspot erosion^5,33,34^.

We found positive selection at the three DNA-binding codons, i.e. positions −1, 3, 6 relative to the a-helix (see figure 5). Structural important sites like the zinc ion ligands are conserved by negative selection. We consider the ongoing selective pressure – positive as well as negative – acting on tarsier *PRDM9* as relevant proof for gene integrity being important in the context of a probable temporary loss of function which was subsequently abolished by a compensatory mutation.

**Figure 5:**
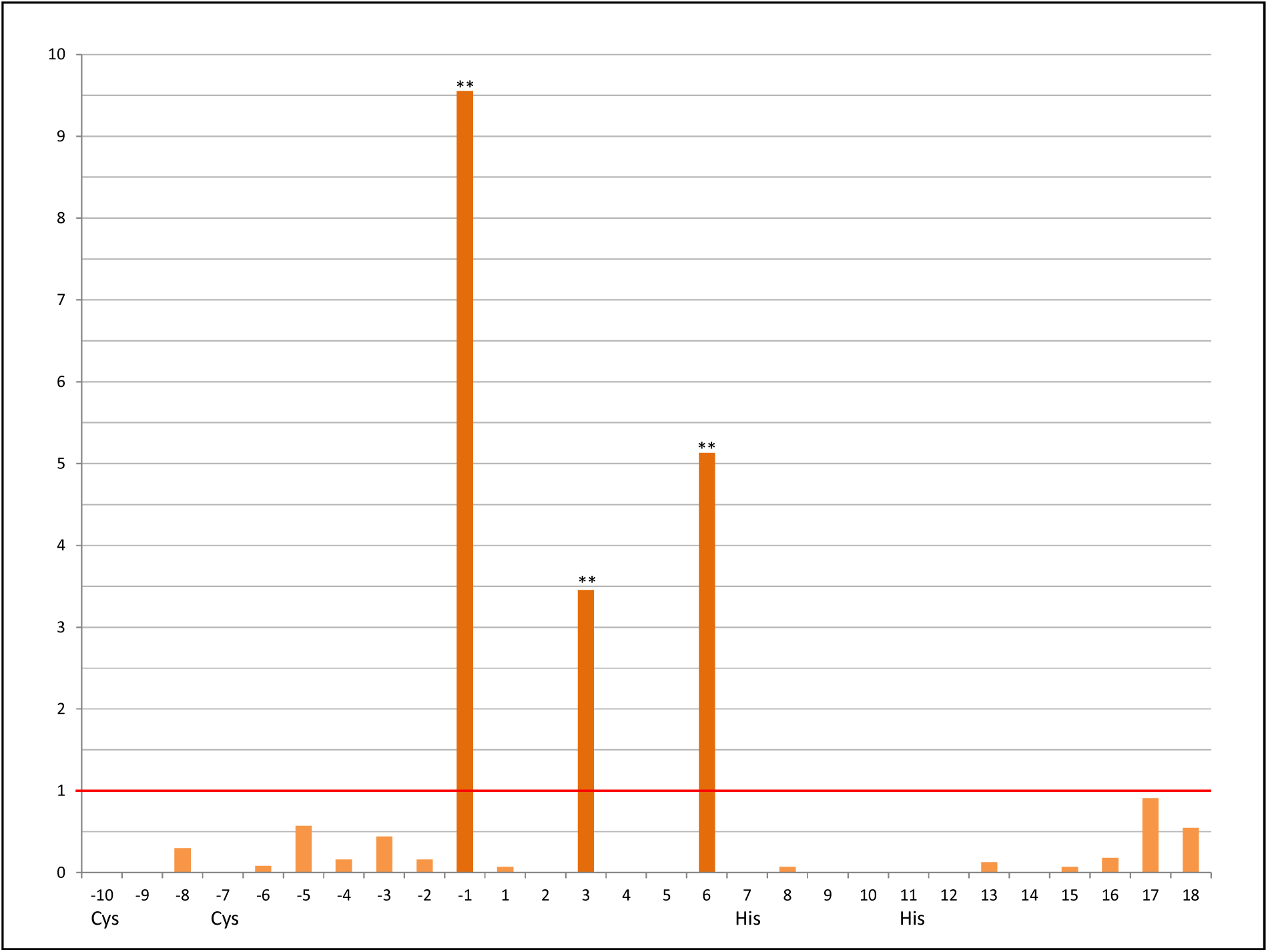
ω-values of C2H2 zinc finger sites in tarsiers. Sites on the x-axis are indicated with their position relative to the beginning of the α-helix. Ligands - cysteine and histidine - are also assigned. The y-axis shows the ω-values with one being the value for neutral selection highlighted by a red line. The three DNA-binding sites (-1, 3, 6) are positively selected witch a p-value < 0.01 after Benjamini-Hochberg correction (**). The other sites show ω-values revealing negative selection with p-values < 0.05 after Benjamini-Hochberg correction (except for the sites −5 to −2, 2, 17 and 18).

## Discussion

With four non-functional zinc fingers at the 5’ end of the tarsier zinc finger array (Fig. 1), degenerated by missing cysteine ligands and frameshifting indels, we detected a completely new *PRDM9* allelic variant in primates and mammals in general. This sequence singularity is also reflected in our inferred phylogenies including first degenerated and all functional C2H2 zinc fingers of the array and clearly designating tarsiers as monophyletic group within haplorhine primates (Fig. 4A and B).

All these sequence autapomorphies, but especially those associated with a temporary loss of allelic integrity (see results) suggest an active role of *PRDM9* in the divergence process between anthropoid and tarsiiform primates or alternatively, along the tarsiid lineage. A possible speciation scenario could be as follows:

In a population of haplorhine or tarsiid ancestors a damaging, frameshifting mutation occurs in one allele (Fig. 2, Fig 6). Heterozygous individuals may be subfertile because gene function is reduced due to one non-functional allele. However, Brick *et al*.^6^ showed that one allele can determine 75% of the hotspots while the other is responsible for 22% [1% of all hotspots are shared and 2% are new in comparison to parental homozygote constellations]. Furthermore, male mice which are heterozygote null for *Prdm9* are not sterile but reproduce at a later stage for the first time and father fewer offspring, both in comparison to the wild-type^35^. Individuals homozygous for the damaged allele are probably infertile due to a considerably reduced or loss of gene function and comparable to *PRDM9* knockout mice where a meiotic arrest at pachytene stage leads to sterility^4^. The second frameshifting mutation restores the functionality of the former erroneous allele (Fig. 2, Fig. 6) and is therefore positively selected and fixated^36,37^. Two of the now three possible allelic variants in homozygotes are functional (Fig. 6), the one with two original or wild-type alleles and the other with two alleles carrying both indel mutations. By contrast, all heterozygotes shown in figure 6 are possibly less fit or rather subfertile resulting from a decreased recombination activity affected by either one damaged allele or different hotspot usage among the two types of functional *PRDM9* alleles^38^. The novel allele finally get fixed in a precursor of tarsiiform primates but in any case no later than 20 MYA, the time to the most recent common ancestor (MRCA) of the three extant tarsier clades^15,17,19^, all exclusively carrying the new allelic variant. Explanations for the fixation of the mutant allele in an ancestral population of modern tarsiers may include directed and/or random evolutionary processes like biased gene conversion or genetic drift. With regard to the former it has been shown that a newly introduced *PRDM9* allele increased hotspot activity in mice through altered and haplotype-biased DNA-binding, thus counteracting fitness loss during the process of hotspot erosion by allelic gene conversion^33^. The mutant allele could have had a similar hotspot enhancing and therewith beneficial fitness effect in a population of ancient tarsiids or their progenitors. Given that its selective advantage was sufficiently large the novel allele could also have overcome the effects of genetic drift where typically rare beneficial mutations are more likely to go extinct^39,40^. On the other hand, we could suppose that it was this process of random change in genetic composition that maintained the newly arisen allele resulting in the structural uniformity of the *PRDM9* zinc finger domain and facilitating divergence of an ancestral lineage finally leading to modern tarsiers.

**Figure 6:**
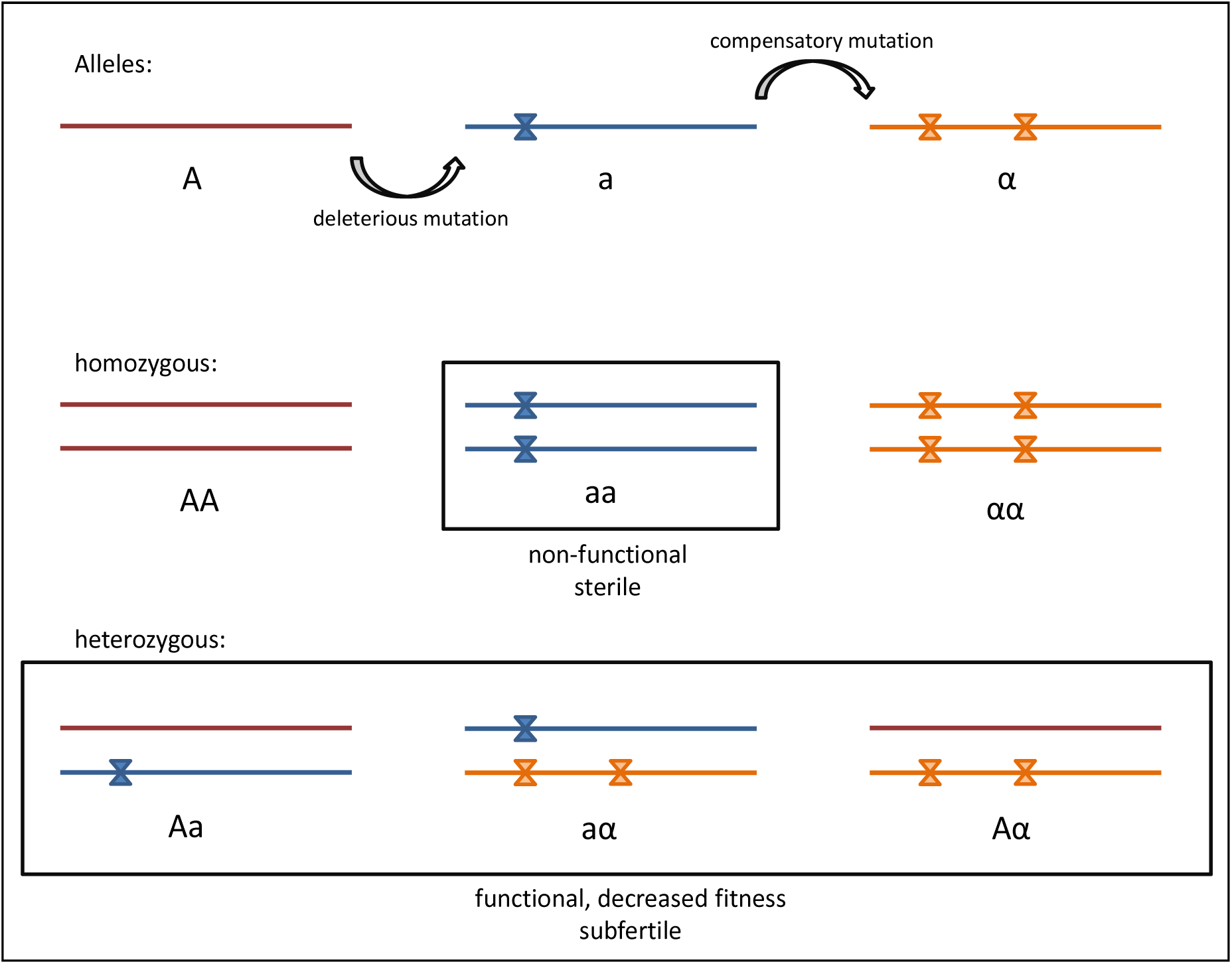
Schematic demonstration of the mutational events in tarsier *PRDM9* zinc finger array. The insertion and deletion create two more alleles, a and α, besides the wild-type allele A. All possible genotypes are shown where all heterozygous are possibly functional but subfertile and two out of three homozygous are functional without constraints and the third is non-functional and sterile, comparable to *Prdm9* knockout mice^6^.

Extinct relatives of living tarsiids branched away from anthropoid primates very shortly after the haplorhine-strepsirhine split^9,17,41^. Depending on the data, either paleontological or molecular, first primates evolved sometime between 55-87 MYA^9,13,17,42^. Within this period two drastic climate changes occurred and both induced mass extinctions^41,43,44^ creating conditions promoting adaptive radiation^41^. Proto-tarsiiforms could have survived or emerged from these evolutionary bottlenecks possibly also because *PRDM9* produced new recombination landscapes allowing allele combinations more favourable or adaptive to the changing environmental conditions and/or newly vacant ecological niches. Another extinction event associated with Eocene-Oligocene cooling^45^ largely decreased primate diversity with the disappearance of omomyids, a controversial fossil tarsiiform primate^46‒48^. This event presumably caused a geographic shift in tarsier distribution from mainland to insular Southeast Asia^49,50^ exposing tarsiids of modern aspects to another population bottleneck and thus providing a further option for the fixation of the novel tarsier-specific *PRDM9* allele. A heightened relevance of genetic drift in tarsier molecular evolution was already discussed^51^, as extant tarsiers, and especially those endemic to the Indonesian island of Sulawesi, have been subject to significant climatic and tectonic change^52,53^ that triggered allopatric speciation in the Malay Archipelago since the Miocene^15, 16, 18, 19, 53‒55^.

*PRDM9* variation in alleles, zinc finger motifs and especially at the key codon positions reflect the different levels of evolutionary independence of the three major tarsier clades (Western, Philippine, and Sulawesi tarsiers). Like it has been observed before, 5´-most and 3´-most C2H2 zinc fingers are more conserved or even identical within or at least between closely related species^38,56,57^ even if positive selected sites are factored in. In tarsiers this appears to apply mainly to species endemic to the same biogeographic region, but is also fairly reflected in lineage-specific constraints on 5´-most C2H2 zinc fingers of Sulawesi tarsiers^19^. Considering the three DNA binding amino acids at internal zinc fingers of Sulawesi tarsiers we found several duplicated zinc finger motifs in each species/subspecies indicating their recent origin^58^ for most populations being estimated at less than 500,000 years ago^19^. Particularly striking are the blocks of zinc finger motifs in northern Sulawesi tarsier populations (Fig. 3). The order of their occurrence very plastically depicts how these populations very recently evolved from a single common ancestor^19^.

Despite the high population specificity of *PRDM9* alleles, we found no clear indication for reproductive isolation by interallelic incompatibilities between parapatric species, since array structure altered even at individual level. The considerable amount of zinc finger motifs shared among Sulawesi tarsiers let us rather suggest that distinct taxa are still hybridizing, as has been shown in previous studies^16,19^. However, the phylogenetic structure of 5´-most zinc fingers is fairly consistent with the two major tarsier lineages on Sulawesi and could therefore be indicative for a PRDM9-mediated post-zygotic isolating barrier among phylogenetic clades.

In conclusion, the high mutation rate of the *PRDM9* zinc finger domain^3,7,59^ and multiple events of genetic drift produced an enormous diversity of *PRDM9* alleles in tarsiers. Nevertheless, inferring reproductive cohesions from *PRDM9* allelic variation seems only to apply for deeper phylogenetic splits in *Tarsius*, presumably also due to current and/or historical hybridization in recently evolved tarsier populations. More intriguing, however, is our discovery of a hitherto unknown and, in addition, tarsier-specific *PRDM9* allele that arose from two indel mutations and played a (perhaps decisive) role in the speciation of early haplorhine primates.

## Material and Methods

### Sample set

The sampling of 23 individuals comprises 21 specimens of nine Sulawesi tarsier populations (1-4 individuals/population, Fig. 7) and two non-Sulawesi tarsiers (*Tarsius bancanus* and *T. syrichta*). Samples from Sulawesi were obtained from previous studies^16,19^, while the Western and the Philippine specimen were provided by Y. Rumpler (Les Hôpitaux Universitaires de Strasbourg, France) and J. Brosius (University of Muenster, Germany), respectively. Whole genome amplifications (WGA) of each sample (40-80 ng/μl) were used as a template in PCR to amplify the exonic region of the *PRDM9* gene containing the zinc finger domain.

**Figure 7:**
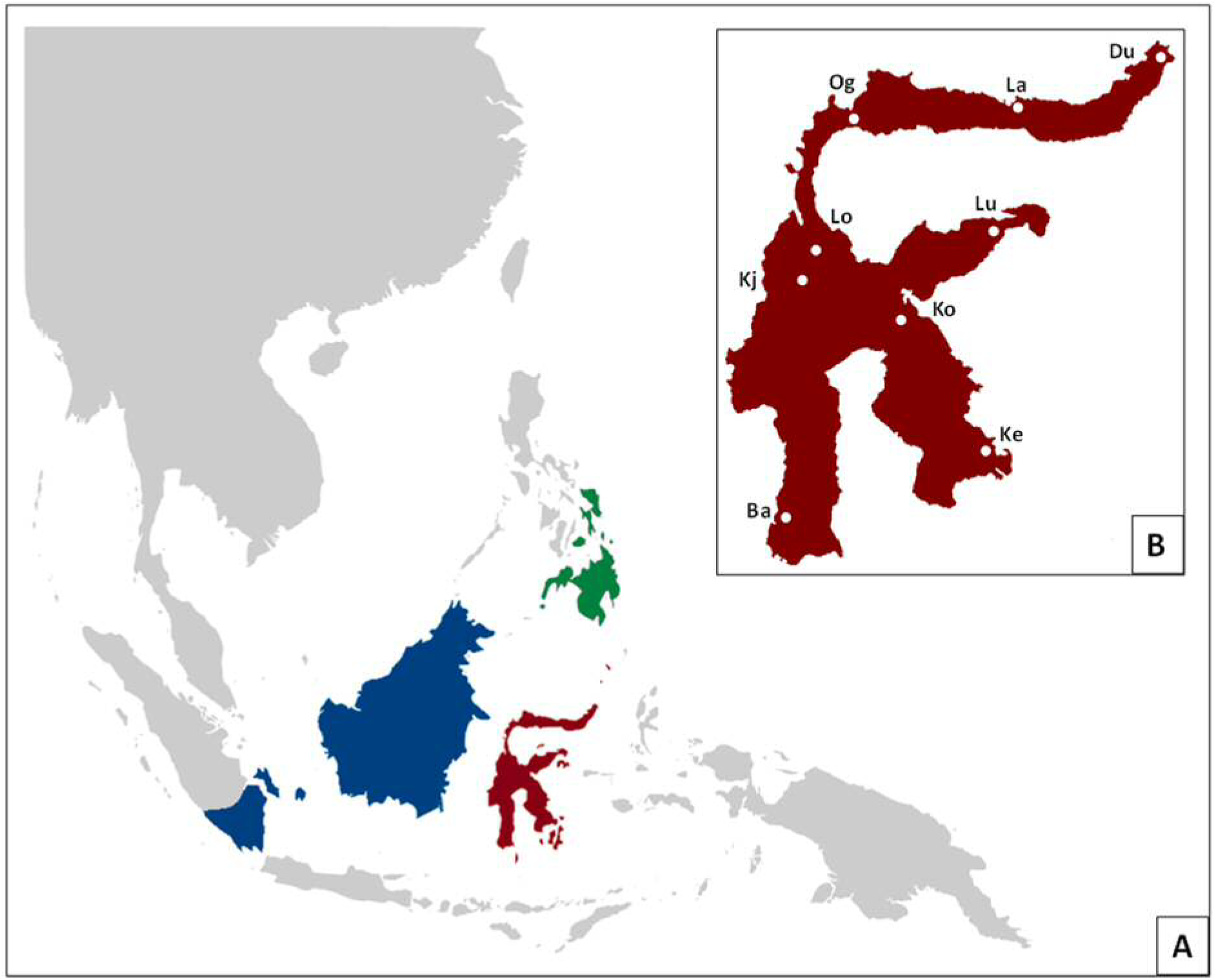
Maps of Southeast Asia and Sulawesi. [A] shows the region of Southeast Asia with ranges of *Tarsius bancanus* in blue, of *Tarsius syrichta* in green and of Sulawesi tarsiers in red. [B] is a close-up of Sulawesi with sample sites highlighted by white dots. Abbreviations: Du, Duasaudara; La, Labanu; Og, Ogatemuku; Lu, Luwuk; Lo, Laone; Kj, Koja; Ko, Korosule; Ba, Bantimurung; Ke, Kendari.

### PCR and Sequencing

We PCR-amplified and sequenced the exon which encodes the zinc finger domain of the *PRDM9* gene in three parts: the 5’ and 3’ flanking regions and the repetitive elements in between (see table 2).

**Table 2:**
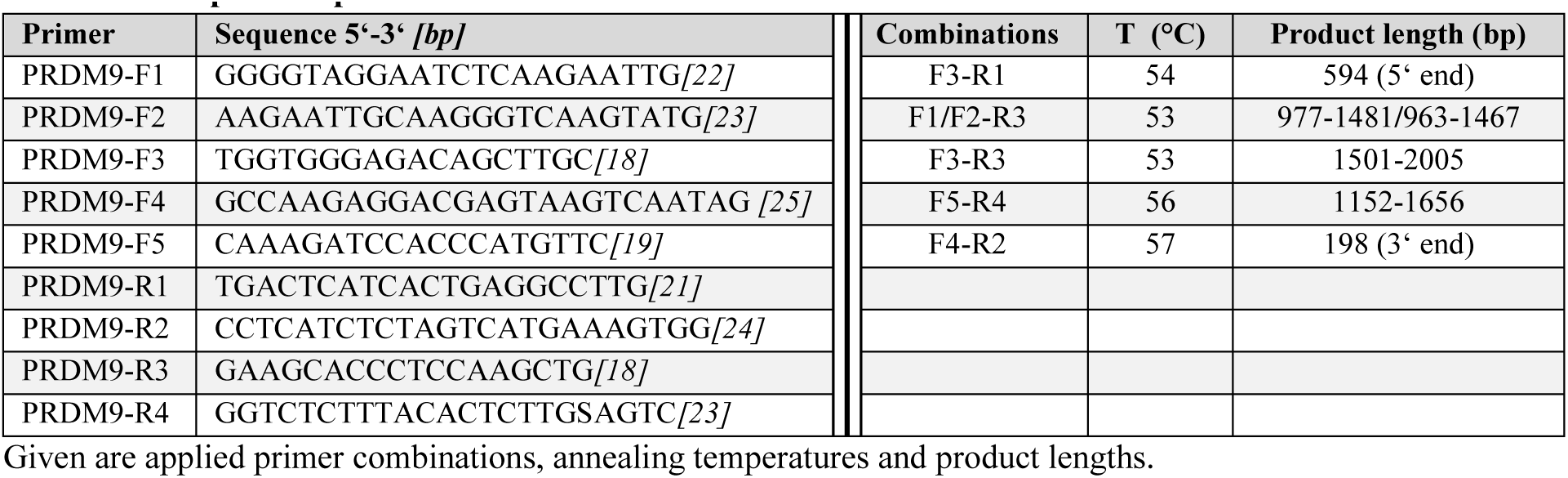
PCR primer specifications.

To reduce amplification of unspecific products we conducted wax-mediated hotstart PCR. Each PCR-reaction contained 30 μl with final concentrations of 200 dNTPs, 2.5 units *Taq* DNA polymerase, 1x PCR buffer including 1.5 mM MgCl_2_ (Qiagen Taq PCR Core Kit), and 0.33 pM per primer. PCRs were run under the following conditions: 3 minutes of initial template denaturation at 94 °C was followed by 35 cycles of denaturation (40 sec at 94 °C), primer annealing (1 min at primer specific temperatures) and DNA elongation (1 – 1.5 min at 72 °C). A final elongation step of 5 minutes at 72 °C finished the PCR. Product sizes were estimated on ethidium bromide stained 1.5 % agarose gels together with O’RangeRuler 100 bp DNA Ladder and GeneRuler 100 bp Plus DNA Ladder (both Fermentas/Thermo Scientific).

In general PCR products were enzyme purified before sequencing. In some individuals PCR reactions yielded multiple different sized products, either because of size variation between the two alleles or unspecific amplicons. The separation of the two alleles or the isolation of the product from non-specific sequences was done by gel extraction with the QIAquick® Gel Extraction Kit (Qiagen). Purified PCR products were sequenced on both strands using the Big Dye Terminator v3.1 Cycle Sequencing Kit (Applied Biosystems) and the corresponding PCR primer pair. After an SDS/heat treatment for elimination of unincorporated dye terminators sequences were processed on a 3130 XL genetic analyzer (Applied Biosystems).

For individuals with ambiguous sites present in their DNA sequences PCR products were purified by ethanol precipitation, ligated into a plasmid vector (pGem®-T Vector System I, Promega) and transformed into One Shot® TOP10 Chemically Competent E. coli cells (Invitrogen). Where gel extraction failed to separate alleles of different lengths sequences were also isolated by cloning.

At least six positive clones, three of each allele, were PCR amplified and sequenced. Colony PCR was carried out in 20 μl reaction volumes according to standard protocols using the Taq PCR Core Kit (Qiagen) and vector-specific primers. For long amplicons where sequencing read length was not sufficient we used the internal primer PRDM9-F2 (see table 2) to get the middle portion of the target sequence.

### Sequence Analyses

Raw sequences were edited and consensus sequences were generated in BioEdit 7.1.3.0^60^. To reveal the structure and integrity of the tarsier PRDM9 gene we created alignments in BioEdit with the following approach. We compared 5´-terminal sequences including the degenerated zinc fingers in front of the C2H2 array 1) within *Tarsius*, and 2) between tarsiers, anthropoids *(Homo sapiens*, ENST00000296682; *Pan troglodytes*, GU166820.1) and rodents *(Rattus norvegicus*, ENSRN0T00000066370; *Mus musculus*, ENSMUST00000167994). We further examined the zinc finger domain by comparison of isolated zinc fingers within and between taxa of *Tarsius*, anthropoids and rodents. Structural and functional properties of tarsier *PRDM9*, in particular with regard to the type of zinc finger motifs, was also validated by peptide sequences obtained with EMBOSS Transeq v. 6.3.1^61,62^.

### Zinc finger Phylogeny

We generated two tree topologies, one including all degenerated first zinc fingers and one including both, functional C2H2-zinc fingers and degenerated first zinc fingers of the array. Haplorhine taxa represented in each data set comprised *Tarsius* (this study), *Homo sapiens* (ENST00000296682) and several other anthropoid primates, whose sequence data were adopted from Schwartz *et al*.^*1*^*. Microcebus murinus* served as outgroup. Corresponding sequence information was extracted from the Ensembl data base (assembly micMur1, database version 77) and by using BLAST+2.2.28^63^. The zinc fingers were isolated with the first codon for cysteine as starting point. All except 5´-most degenerated and several 3´-most zinc fingers had a length of 84 bp. Lacking the first cysteine-triplet 5´-most degenerated zinc fingers start with the second codon for cysteine, and therefore being nine bp shorter. 3’-most zinc fingers can lack base pairs when interrupted by a stop codon, but they contain at least both cysteine and histidine ligands. Like Schwartz *et al*^7^. we excluded the four binding triplets. To reduce the calculation time we removed identical zinc fingers within genera with the Fabox DNAcollapser^64^. Models of nucleotide substitution were determined with jModelTest 2.1.6^65,66^. Phylogenetic trees were generated with PhyML 3.0^65^ including an approximate likelihood ratio test (aLRT) and SH-like supports. The degenerated zinc finger of *Microcebus murinus* served as outgroup in both tree topologies. Nodes with support values below 0.5 were collapsed in TreeGraph 2.2.0-407 beta^67^. The final phylogenetic trees were visualized in FigTree 1.3.1^68^.

### Detecting selective pressure

All functional C2H2 zinc fingers (84 bp) were tested for selective pressure on codon sites. We inferred a phylogenetic tree based on all non-identical sequences determined by Fabox DNAcollapser^64^ using PhyML 3.0 [settings other than default: K80, bootstrap: 100]^65^. The best fitting substitution model was selected using jModelTest 2.1.6^65,66^. Tests of selection were performed by SLR^69^ using default settings.

## Acknowledgments

This work was supported by grants (ZI568/6-1 and ZI568/6-2) from the Deutsche Forschungsgemeinschaft (DFG, http://www.dfg.de) allocated to H.Z.

